# sRNAanno --- a database repository of uniformly-annotated small RNAs in plants

**DOI:** 10.1101/771121

**Authors:** Chengjie Chen, Junting Feng, Bo Liu, Jiawei Li, Lei Feng, Xiaoling Yu, Jixian Zhai, Blake C. Meyers, Rui Xia

## Abstract

Small RNAs (sRNAs) are essential regulatory molecules, including three mayor classes in plants, microRNAs (miRNAs), phased small interfering RNAs (phased siRNAs or phasiRNAs), and heterochromatic siRNAs (hc-siRNAs). Except miRNAs, the other two classes are not well-annotated and collected in public databases for most sequenced plant genomes. We performed comprehensive sRNA annotation for 138 plant species, which have fully sequenced genomes and public next-generation-sequencing (NGS) sRNA data available. The results are available via an online repository called sRNAanno (www.plantsRNAs.org). Compared to plant miRNAs deposited in miRBase, we obtained much more miRNAs, which are more complete and reliable because of consistent and high-stringent criteria used in our miRNA annotation. sRNAanno also provides free access to genomic information for >16,000 *PHAS* loci and >21,000,000 hc-siRNA loci annotated from these 138 plants. On the basis of Integrative Genomics Viewer (IGV), we developed a visualization tool for browsing NGS sRNA data (IGV-sRNA), which have been integrated a series of new functions compatible to specific sRNA features. To make sRNA annotation an easy task, sRNAanno also provides free service of sRNA annotation to the community. In summary, sRNAanno and IGV-sRNA are great resources to facilitate the genomic and genetic research of plant small RNAs.

## Introduction

Small RNAs (sRNAs) are essential regulatory molecules in plants. With the fast development of deep sequencing technologies and bioinformatics, sRNAs have been characterized in more and more plant species, leading to the generation of huge amount of next generation sequencing (NGS) data. A large number of raw NGS data of sRNAs have been deposited in public database, such as Gene Expression Omnibus (GEO) and European Nucleotide Archive (ENA). MicroRNAs (miRNAs) are the most well-studied class of sRNAs in plants. To date, miRBase is the primary repository and online database for annotated miRNAs (Kozomara et al., 2019). As a routine practice in the research community, the annotated miRNAs of a species are required to deposit into miRBase before publication, i.e., author submission is the primary data source of the database. This process makes it hard to maintain a high quality of annotated miRNAs deposited in miRBase because of the variable stringency of the criteria, controlled by the submitting authors who are responsible for miRNA annotation. In other words, rather than the developers or maintainers of miRBase, quality control is more reliant on the authors, reviewers and editors, who likely have different understanding of the criteria of miRNA annotation. Therefore, the variable reliability of annotated miRNAs in miRBase is a great concern to the community (Axtell and Meyers, 2018).

In addition to miRNAs, there are other types of sRNAs in plants, including phased small interfering RNAs (phased siRNAs or phasiRNAs) and heterochromatic siRNAs (hc-siRNAs). phasiRNAs have recently been emerging as critical regulatory molecules in almost all aspects of growth and development in plants (Fei et al., 2013). They are widely present in plants from algae to angiosperms. However, compared to miRNAs, they are much less studied and are not well annotated in most of the full genome sequenced plants. To date, there is no public database collecting the annotation information of plant phasiRNAs, hindering the application of already annotated phasiRNA information. Heterochromatic siRNAs are the most abundant class of sRNAs in plants, and they usually play roles related to DNA methylation, which is a process important for transcriptional regulation. Although we know its functional importance, thorough annotation of hc-siRNA-generating genomic regions (hc-siRNA loci) is always lacking for most plant genomes.

In this study, we conducted extensive sRNA annotation for 138 plant species with full-genome sequenced and at least one sRNA deep-sequencing dataset available in public databases. The annotation includes all these three sRNA classes, miRNAs, phasiRNAs and hc-siRNAs. To achieve a high confidence for miRNA annotation, we applied a set of uniform criteria adopted from the recently updated rules for plant miRNA annotation (Axtell and Meyers, 2018). For phasiRNA annotation, a p-value based approach established by our group was used for annotation of loci generating 21-nt phasiRNAs (21-*PHAS*) or 24-nt phasiRNAs (24-*PHAS*) (Xia et al., 2015, 2019). We also developed an algorithm based on sequence repetitiveness for the accurate annotation of loci generating hc-siRNAs, given their primary feature of generation from repetitive genomic regions. In total, we annotated 23,908 miRNA hairpins or precursors, 16,797 *PHAS* loci (15,121 21-*PHAS* and 1,676 24-*PHAS*) and 21,841,048 hc-siRNA loci. All those results were deposited in an online database of sRNA annotation (sRNAanno) for open access. In addition, to facilitate the visualization of annotation results for manual check, which is a critical step to maintain the high quality of annotation, we developed a tool for the exploration of sRNA data (IGV-sRNA), in which new functions specifically designed for sRNA features were integrated into the Integrative Genomics Viewer (IGV)(Thorvaldsdóttir et al., 2013).

## Database content

### sRNAanno database

Small RNA annotation for three major sRNA classes was performed for 138 plant species, and an online database (sRNAanno, www.plantsRNAs.org) was constructed to store all the annotation results for easy and quick public access. There are three major functions in sRNAanno, including BROWSE of annotation results, SEARCH for certain information and RESOURCE for data sharing (Fig. 1A). In the BROWSE page, users can select single or several species from a large phylogenetic tree and download corresponding small RNA annotation files in GFF3 format. The SEARCH function includes miRNA searches by either miRNA name or sequence and *PHAS* sequence comparison using the BLAST function. In addition, we provide quick access to download the tool IGV-sRNA, which is designed for the exploration of sRNA data (described in more detail below).

**Fig. 1.**
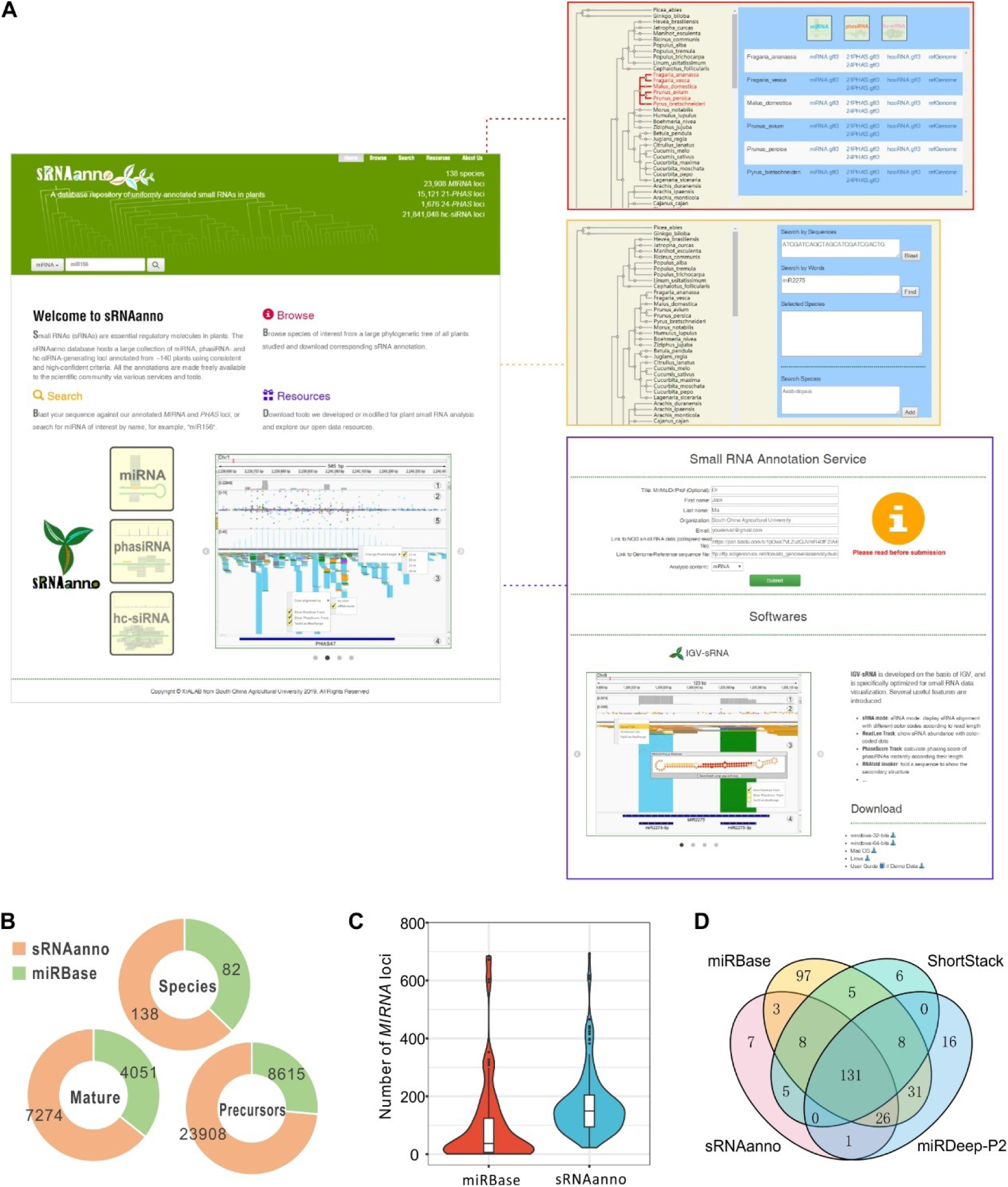
Overview of the sRNAanno database. A. Screenshots of the HOME page, and subpages for three main functions: BROWSE, SEARCH, and RESOURCE. B. Comparison between miRBase and sRNAanno on numbers of plant species and *MIRNA* loci annotated. C. Number of *MIRNA* loci annotated in each species in miRBase and sRNAanno. D. Comparison of the miRNA annotation results of sRNAanno for Arabidopsis with those from three other sources/pipelines, miRBase, shortstack and miRDeep-P2.

### miRNA annotation

In this study, we aimed to conduct a genome-wide annotation of plant miRNAs using a set of uniform and well-established criteria (Fig. S1), as discussed in (Meyers et al., 2008; Axtell and Meyers, 2018). To this end, we downloaded the genome sequences for almost all species for which genomes and their sRNA NGS data are both available from public databases (such as ENA and GEO). We found 138 plants with corresponding sRNA data available. In total, there are 1,586 small RNA sequencing datasets, with most from well-studied model plants, like Arabidopsis and rice. We performed miRNA annotation for all these species, and we obtained 23,908 annotated hairpin precursors, encoding 7,274 unique mature miRNA sequences (Fig. 1B). Compared to the latest release of miRBase (v22), which contains 8,615 annotated hairpin precursors from 82 plant species with 4,051 mature miRNA sequences (Fig. 1B), our annotation yielded more results, in terms of not only the number of species annotated (a 1.68 fold increase), but also the number of *MIRNA* precursors (a 2.78 fold increase) (Fig. 1B). In plants, there are ∼24 miRNA families prevalently present in angiosperms (Chen et al., 2018). To assess the completeness of miRNA annotation of a species, we compared the number of conserved miRNA families in species which have data in both sRNAanno and miRBase. We found that almost all of the 45 species analyzed have a more complete list of conserved miRNAs in sRNAanno, while conserved miRNAs in 14 species are obviously incomplete in miRBase (v22) (Fig. S2). Moreover, in terms of the number of *MIRNA* precursors annotated from a single organism, most species have fewer *MIRNAs* in miRBase (<100) compared to sRNAanno (100-200, Fig. 1C). Therefore, sRNAanno has more complete miRNA data than miRBase (v22). When compared with miRNAs deposited in miRBase or identified by other tools (miRDeep-P2 and ShortStack) within a species (using Arabidopsis and rice as examples), we found that the majority of miRNAs annotated in sRNAanno were identified as well by at least one of the other tools ((Kuang et al., 2019; Axtell, 2013b), Fig. 1C and Fig. S3); only a few of them were unique to sRNAanno. Taken together, we contend that our miRNA annotation is of high stringency and high confidence.

### *PHAS* locus annotation

Phased siRNAs (phasiRNAs) are another major class of sRNAs found in plants, and these are universally present in all plants, mainly as the trans-acting siRNA (tasiRNA) subgroup (Xia et al., 2017). This group characterized by the “phasing pattern” of sRNAs-approximately a head-to-tail arrangement. To date, unlike miRNAs, there is no database collecting the identified or reported *PHAS* genes or genomic loci, although phasiRNAs have been profiled in a large number of plants (Fei et al., 2013). Therefore, we performed exhaustive *PHAS* profiling for the 138 plant species with at least one sequenced sRNA library. We used a well-developed p-value-based protocol to perform *PHAS* analysis ((Xia et al., 2013, 2019), Fig. S4). The cutoff of the p-value was set to 1e-10. For the analysis of 24-*PHAS* loci, we added an additional filter to remove repetitive sequences which usually give rise to abundant 24-nt hc-siRNAs. In total, we identified 15,121 21-*PHAS* loci generating 21-nt phasiRNAs, and 1,676 24-*PHAS* loci producing 24-nt phasiRNAs. In general, the number of 21-*PHAS* loci was substantially greater than that of 24-*PHAS* loci within a species (Fig. 2A). Both types of *PHAS* loci are not evenly present across species (Fig. 2A), perhaps because of the sampling for sRNA sequencing (tissue, stage, etc.) and the sequencing technology (sequencing platforms, sequencing depth, etc.). Another factor accounting for this uneven distribution of *PHAS* loci is likely the intrinsic genomic difference among plant species, for instances, plants from certain families, like Brassicaceae (including the model plant Arabidopsis), and Cucurbitaceae, consistently yield fewer *PHAS* loci compared to other species (Fig. 2A). As reported before, 24-*PHAS* are more widely existed in monocots, but dispersed in eudicots without noticeable pattern ((Xia et al., 2019), Fig. 2A)

**Fig. 2.**
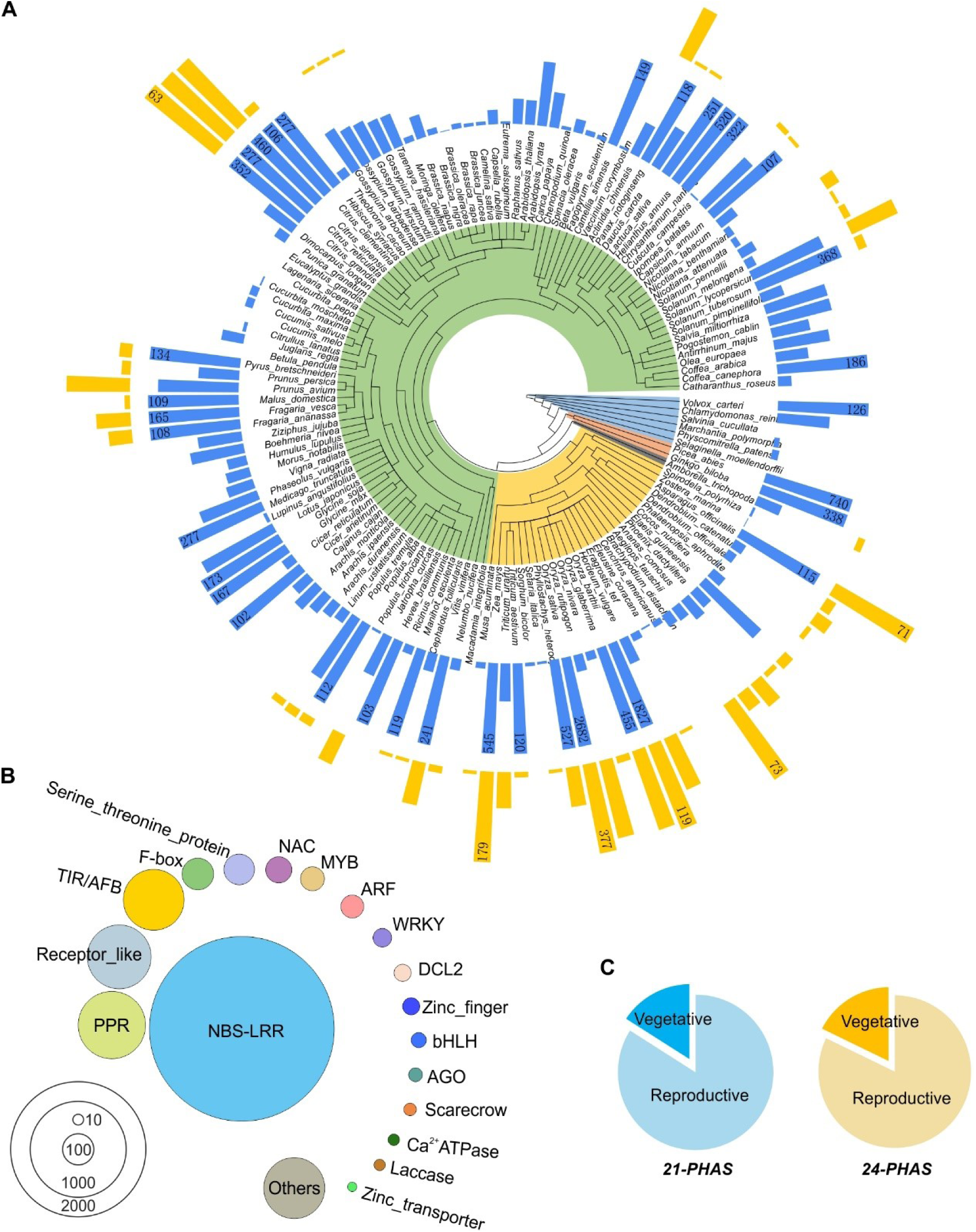
Statistics of annotated *PHAS* loci in sRNAanno. A. Numbers of 21-(inner circle, blue) and 24-(outer circle, yellow) *PHAS* loci annotated in each species. All species are ordered according to the phylogenetic tree of APG IV (Angiosperm Phylogeny Group IV). B. Functional classification of protein-coding 21-*PHAS* loci. Circle sizes are scaled by the number of 21-*PHAS* loci in a given gene group. C. Tissue specific enrichment of 21- and 24-*PHAS* loci.

Protein coding genes are a rich source of phasiRNAs. After the functional annotation of 21-*PHAS* loci and assessment of their protein-coding capacity, we found that a large number of gene families produce profuse phasiRNAs, especially for gene families of *NBS-LRR, PPR, Receptor-like kinase*, etc. (Fig. 2B). In particular, many transcription factor genes produce phasiRNAs, including *TIR/AFB, F-box, NAC, MYB, ARF, WRKY, Zinc finger*, and *bHLH* (Fig. 2B). In terms of noncoding *PHAS* loci, for both 21-*PHAS* or 24-*PHAS*, most of them were enriched in reproductive tissues (Fig. 2C).

### hc-siRNA locus annotation

hc-siRNAs account for a major part of the plant cellular sRNA population. Currently the genomic loci producing hc-siRNAs have not been well annotated, due to their large number, variability in sRNA abundances, and the complexity of their biogenesis. hc-siRNAs are generated from repeat-related sequence regions, typically transposons, and heterochromatic regions, to direct *in cis* DNA methylation in plants (Axtell, 2013a). As these regions generating hc-siRNA are also an important component of genomic information for a species, we annotated hc-siRNA loci for genomes with sRNA data available according the criteria listed in Fig. S5. Indeed, these hcsiRNA-generating loci are copious in almost every genome (Fig. S6).

### IGV-sRNA for visualization of NGS sRNA data

Currently, all sRNA annotation tools or pipelines have limitations. No matter which tools or pipelines are used for small RNA annotation, there are potential misannotations, especially for miRNA and 24-nt phasiRNAs (Polydore et al., 2018). A manual check of the computationally annotated results of raw genome-mapping data is a suggested practice and a reliable way to minimize misannotations (Xia et al., 2017). Integrative Genomics Viewer (IGV) is a widely used open-source visualization tool for exploring deep sequencing data (Thorvaldsdóttir et al., 2013). It is known for its user-friendly interface and good compatibility to all kinds of genomic data stored in various formats. It is specialized for viewing genome-sequencing or resequencing data or RNAseq data. Due to the high diversity of the biogenesis and function of plant small RNAs, IGV cannot represent all the features of small RNAs as desired, for instance, the length of sRNA reads, which is a critical feature of small RNA biogenesis and function, and the secondary structure of sRNA-generating loci, which is an indispensable feature of *MIRNA* precursor genes.

Therefore, we have developed, on the basis of IGV, a few new functions specifically for the visualization of various characteristics of sRNAs. The modified browser tool was named IGV-sRNA. Its new functions/features include the following:

#### a. Automatic resolution of the genome-mapped file of collapsed sRNA datasets

Different to RNAseq or resequencing DNA data, raw sRNA data are of high sequence redundancy, i.e., a sRNA may be of multiple copies, and will be sequenced multiple times in a library, then giving rise to many identical reads in a sequence file. After sequence collapse, reads of identical sequences will be merged together and each sRNA sequence will be unique in a collapsed dataset, leading to smaller data file size and saving time and computation resources for the following analyses and data exchange. Since IGV was originally developed for the exploration of non-redundant data, such as RNAseq or resequencing DNA data (or other data), it does not support collapsed sRNA files. Every collapsed sRNA will be shown only one time as a single read without displaying its abundance information (Fig. S7). To explore sRNA datasets, we previously decompiled the collapsed sRNA files, which greatly increases the file size, needs more computation resources and slows down the performance of IGV. To address these issues, we incorporated a new feature into the IGV-sRNA to support collapsed sRNA files, which will normally contribute to a 5 to10 fold decrease in the file size. When loaded with the genome-mapped files of collapsed sRNA datasets, the IGV-sRNA will automatically resolve the collapsed reads to show the read abundance of each sRNA sequence, while the original IGV will represent each sRNA only one time in the browser (Fig. S7).

#### b. Display sRNA with different color code according to their length

As the length of sRNAs is an important feature indicating their biogenesis and function, it is useful to know the length when viewing sRNA data. For this purpose, we adopted the color code used in the web browser of MPSS (massively parallel signature sequencing) (Nakano et al., 2006), with the cyan color for 21-nt reads, green for 22-nt, purple for 23-nt, orange for 24-nt, and grey for others (Fig. 3). In the sRNA mode of IGV-sRNA, all the reads can be shown with different colors, which is helpful for quick assessment of sRNA length (Fig. 3).

**Fig 3.**
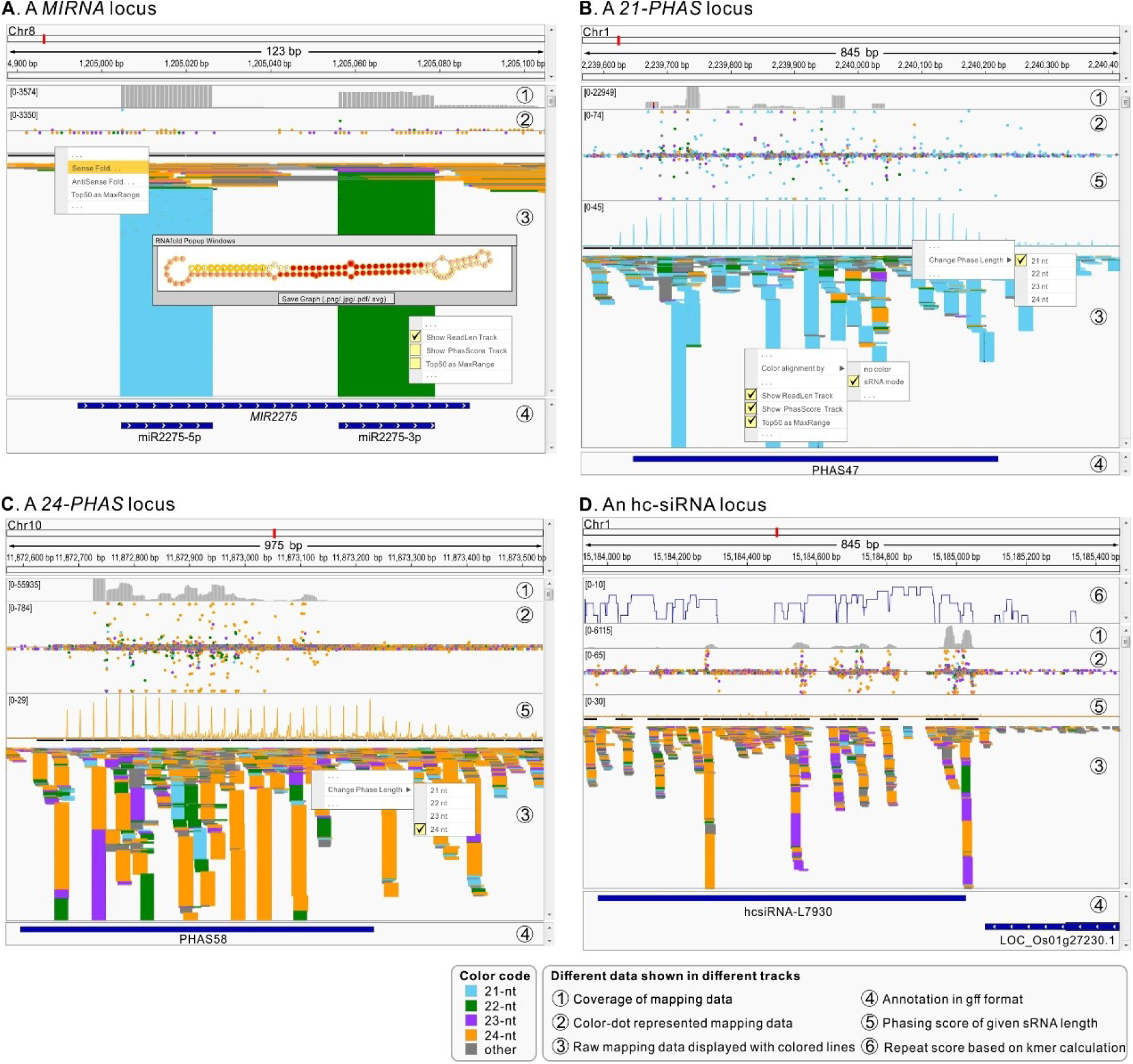
Representative sRNA-generating loci viewed with IGV-sRNA. **A** A representative *MIRNA* locus. Users can fold a sequence to view the secondary structure with coverage information indicated synchronously. The abundance of each sRNA could be showed in color-coded dot plots (track #2), with the cyan color for 21-nt reads, green for 22-nt, purple for 23-nt, orange for 24-nt, and grey for others. All alignments in IGV-sRNA are displayed with different color code according to read length (track #3). **B** and **C** Representative 21-*PHAS* locus and 24-*PHAS* locus. Phasing score of phasiRNAs can be calculated instantly according their length and display by line plot (track #5). **D** A representative hc-siRNA locus. 23-24 nt sRNA are enriched in this region with a high repeat-score (track #6).

#### c. Show sRNA abundance with color-coded dots

The original IGV cannot properly show the abundance of each sRNA sequence, and it can only be approximately assessed by the track of coverage (sequence depth at each genomic position). To obtain accurate abundance information while browsing sRNA data, we adopted the sRNA dot plot using the same color code to show sRNA length, with the leftmost nucleotide to show the genomic position (Fig. 3). In some cases, a single or few extremely abundant sRNAs will cause other less abundant sRNAs distributed close to the X axis. To avoid this, we provide an option to set the maximum value of Y axis to “Top-50 read abundance” (the abundance ranked 50th for all the sRNAs in the current viewing window), with these sRNAs of abundance larger than the maximum value display in triangles (instead of dots) close to the top or bottom of the data track.

#### d. Fold a sequence to show the secondary structure with coverage information indicated synchronously

A leading feature of a miRNA is that its precursor gene possesses a good stem-loop (hairpin) structure. Therefore, to facilitate the manual evaluation of annotated miRNA candidates, we developed a new function in IGV-sRNA to allow instant folding of a sequence to assess its secondary structure, as shown in Fig. 3A. Moreover, the coverage information of each nucleotide can be represented simultaneously with color intensity (Fig. 3A).

#### e. Calculate the phasing score of phasiRNAs instantly according to their length and display using line plot

The phasing pattern is the determining feature of phasiRNAs, and the phasing score is an important metric with which to evaluate the strength of phasing pattern. To assist the quick evaluation of phasing pattern, we developed a new function to do the calculation of phasing score instantly according to different sRNA length in IGV-sRNA (Fig. 3B). The phasing score is shown in a separate track, by line plot, which is also color-coded with the same color scheme mentioned above (Fig. 3B and 3C). Whether a sRNA-generating locus is a *PHAS* locus or not can be quickly evaluated by the data track of phasing score.

#### f. Show pre-calculated repeat score of a genome

As aforementioned, hc-siRNAs reprent the most abundant class of sRNA. They are mostly derived from repetitive sequence regions, like transposons. To quickly assess the repetitiveness of a sequence, we can load the pre-calculated data of repeat score into IGV-sRNA to distinguish hc-siRNAs with other types of 24-nt siRNAs, for instance, 24-nt phasiRNAs and 24-nt siRNAs derived from long inverted-repeat regions (Fig. 3D).

### sRNA annotation service

Small RNA data analysis using various bioinformatics tools or pipelines relying on programming and command-line environment is challenging and time-consuming for most wet-lab biologists. To make sRNA annotation an easy task, we are providing free service of sRNA annotation in sRNAanno. Users can upload the sRNA NGS data and corresponding genome/reference sequence file to an accessible online repository (like an FTP site) and submit the information of download links to these files to sRNAanno in the “Resources” page. Upon receiving the information, we will download the data files, perform the sRNA annotation, and return the annotation results to the users by email. For this service, we will keep high-confidential of user’s data and results, and will not, in any circumstance, use them or release to public without user’s permission.

## Discussion

The new database repository of plant small RNAs described here, sRNAanno, is a repository of major types of sRNAs for nearly 140 plant genomes. This extensive annotation was achieved by analyzing ∼1,600 sRNA datasets, using well-established computation pipelines with reliable and highly stringent criteria. The sRNAanno includes miRNA annotation from ∼33% more plant species than miRBase, the main and most popular miRNA hub. On the other hand, all the miRNAs in sRNAanno were annotated via an identical process with consistent criteria, in comparison to the variable annotation criteria used for the miRNAs in miRBase, whose annotation was conducted by different research groups with various tools (Nakano et al., 2006). Generally, we believe the miRNA annotation of sRNAanno is more reliable compared to miRBase. However, there is no golden rule for the annotation of plant miRNAs. Although there are misannotations in miRBase, there may be a certain number of bona fide miRNAs which are possibly missing in sRNAanno. Moreover, miRBase also deposits miRNAs from plant species without sequenced genomes or without public sRNA data available (for which we are unable to perform miRNA annotations). Therefore, sRNAanno will be a good complement, instead of a substitute, to miRBase.

In addition to miRNA, sRNAanno also stores genomic loci generating phasiRNAs or hc-siRNAs. Annotation of these sRNA-generating loci was conducted using high confidence settings according to the widely accepted criteria. In plants, phasiRNAs have been emerging as one of the major types of sRNAs, and their targets function in a broad range of biotic and abiotic processes. For instance, phasiRNAs are profusely produced during the reproductive stage, especially in the monocots and their subgroups of grasses (Xia et al., 2017)(Zhai et al., 2015). In rice there are >2000 *PHAS* loci generating 21-nt phasiRNAs and ∼400 loci 24-nt phasiRNAs (Tamim et al., 2018). Although phasiRNAs, including the well-known tasiRNAs, have been characterized in more and more plant species, there is no public online repository collecting reported or annotated phasiRNAs and provide convenient and quick access to this information. Similarly, hc-siRNAs are widely present in plant cells, and they are well-known for their connection to DNA methylation or other epigenetic modifications, but the majority of plant genomes lack a good annotation of hc-siRNA loci. In this study, we performed a broad annotation of phasiRNAs and hc-siRNAs in plants, and the resulting annotation stored in sRNAanno are a valuable resource to facilitate genomic and genetic research in plants.

Both manual check of annotation results and data representation need efficient exploration of raw NGS data. Due to their specific features, visualization of NGS sRNA data is distinct from RNAseq or other genomic data. The development of IGV-sRNA with a series of new functions ensures an excellent compatibility of IGV to display sRNA data. These specifically designed functions, like showing read length in color scheme, fold of secondary structure, and instant calculation and visualization of phasing score, significantly improve the compatibility of IGV to NGS sRNA data, and make the exploration of sRNA data more efficient and pleasant.

## Conclusions

Thorough annotation of miRNAs, phasiRNAs, and hc-siRNAs was conducted for 138 plant genomes. All the annotation results are of high quality and confidence, and have been deposited in a public database repository sRNAanno (www.plantsRNAs.org) for quick and convenient access. A visualization tool, IGV-sRNA, was developed as well with specific functions for the efficient exploration of NGS sRNA data. All these results or tools will be valuable resources facilitating the research on sRNAs or related areas in plants.

## List of abbreviations

NGS: Next generation sequencing;
sRNAs: small RNAs;
miRNAs: microRNAs;
siRNAs: small interfering RNAs;
phasiRNA: phased siRNAs;
hc-siRNAs: heterochromatic siRNAs;
IGV: Integrative Genomics Viewer.

## Availability of data and material

All the open access data (including all the annotated sRNA results) are available through the sRNAanno database (www.plantsRNAs.org)

## Competing interests

The authors declare that they have no competing interests

## Authors’ contributions

CC and RX designed the study. CC, JF, JL, LF, BL, and XY collected the data and performed the analyses. JZ and BCM helped with the data analysis and database design. CC and RX constructed the database and wrote the manuscript.

## Acknowledgements

This work was funded by the National Key Research and Developmental Program of China (2018YFD1000104). This work was also supported by awards from the National Natural Science Foundation of China (#31872063), the Guangzhou Science and Technology Key Project (201804020063), the Innovation Team Project of the Department of Education of Guangdong Province (2016KCXTD011), and the Key Areas of Science and Technology Planning Project of Guangdong Province (2018B020202011). Group of J.Z. is supported by the Guangdong Innovative and Entrepreneurial Research Team Program (2016ZT06S172). We thank all members of the Xia lab for their help on this project.

---

**Figure S1.**
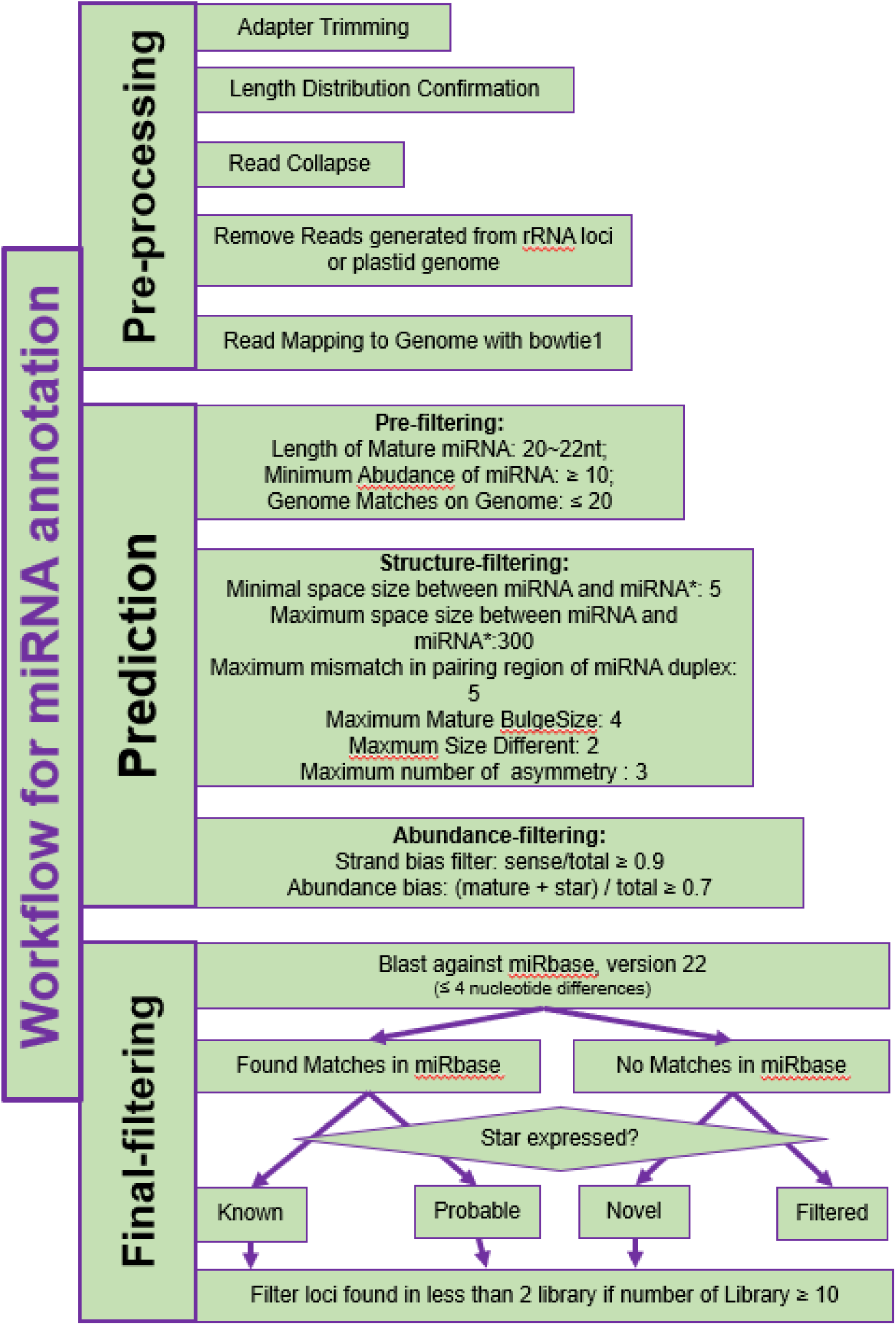
Workflow of miRNA annotation.

**Figure S2.**
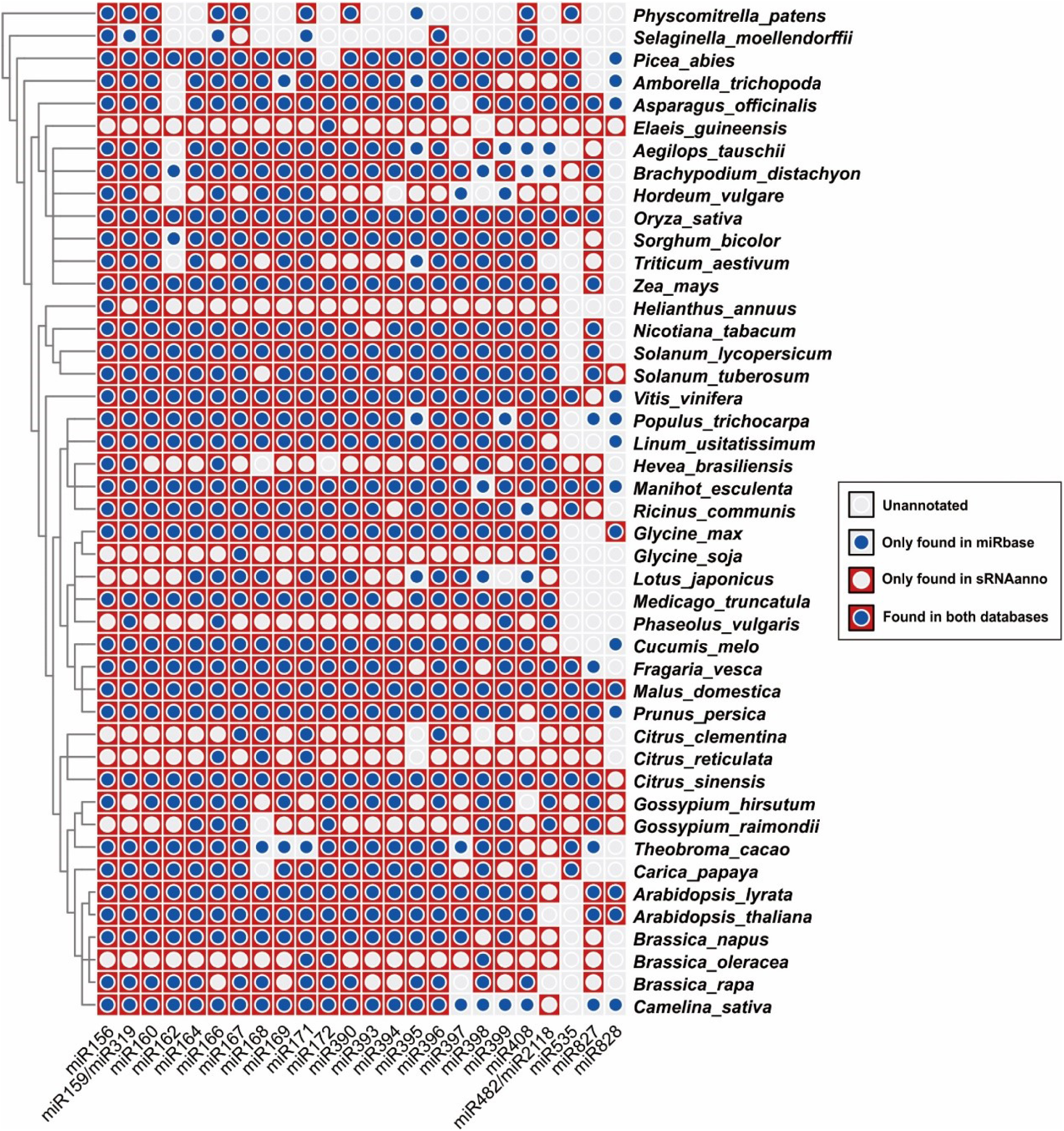
Conserved miRNAs overlapped between sRNAanno and miRbase. Conserved miRNAs of 45 plants were compared between sRNAanno and miRbase. All species are ordered according to the phylogenetic relationship of APG IV. Each cell represents whether a miRNA family (column) is annotated in a species (row) or not.

**Figure S3.**
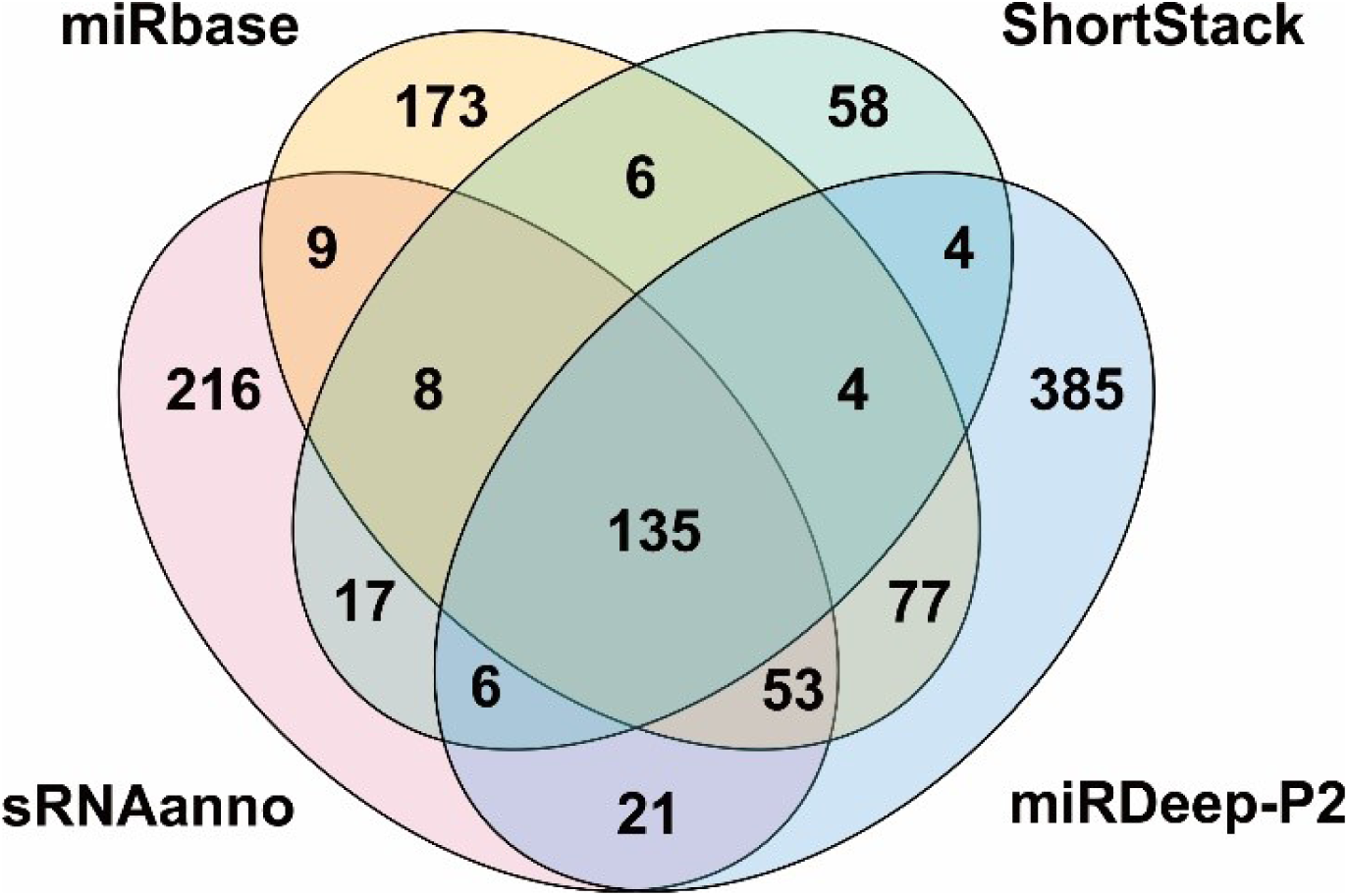
Comparison of miRNA annotation results of rice from miRbase, ShortStack, sRNAanno and miRDeep-P2.

**Figure S4.**
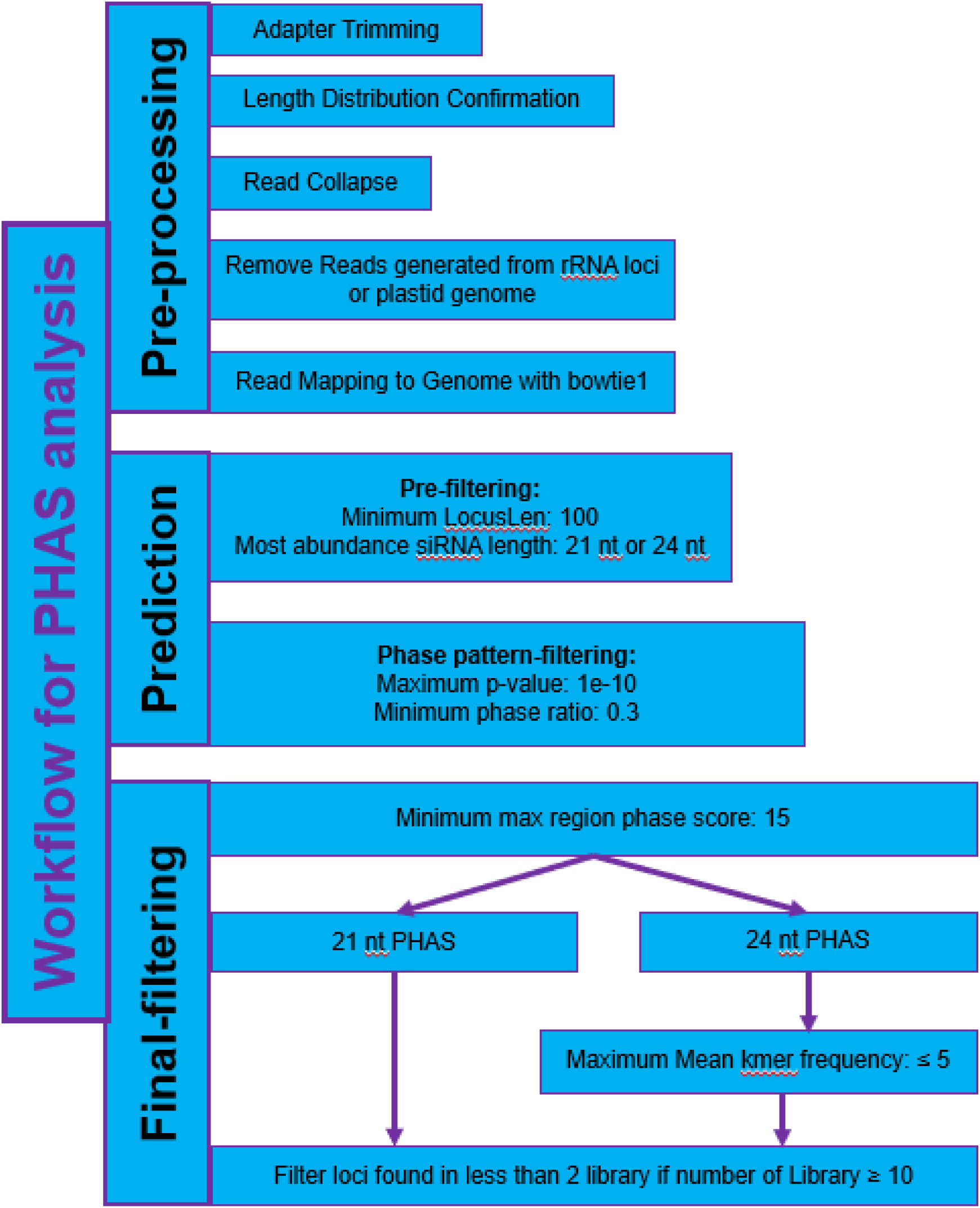
Workflow of *PHAS* analysis.

**Figure S5.**
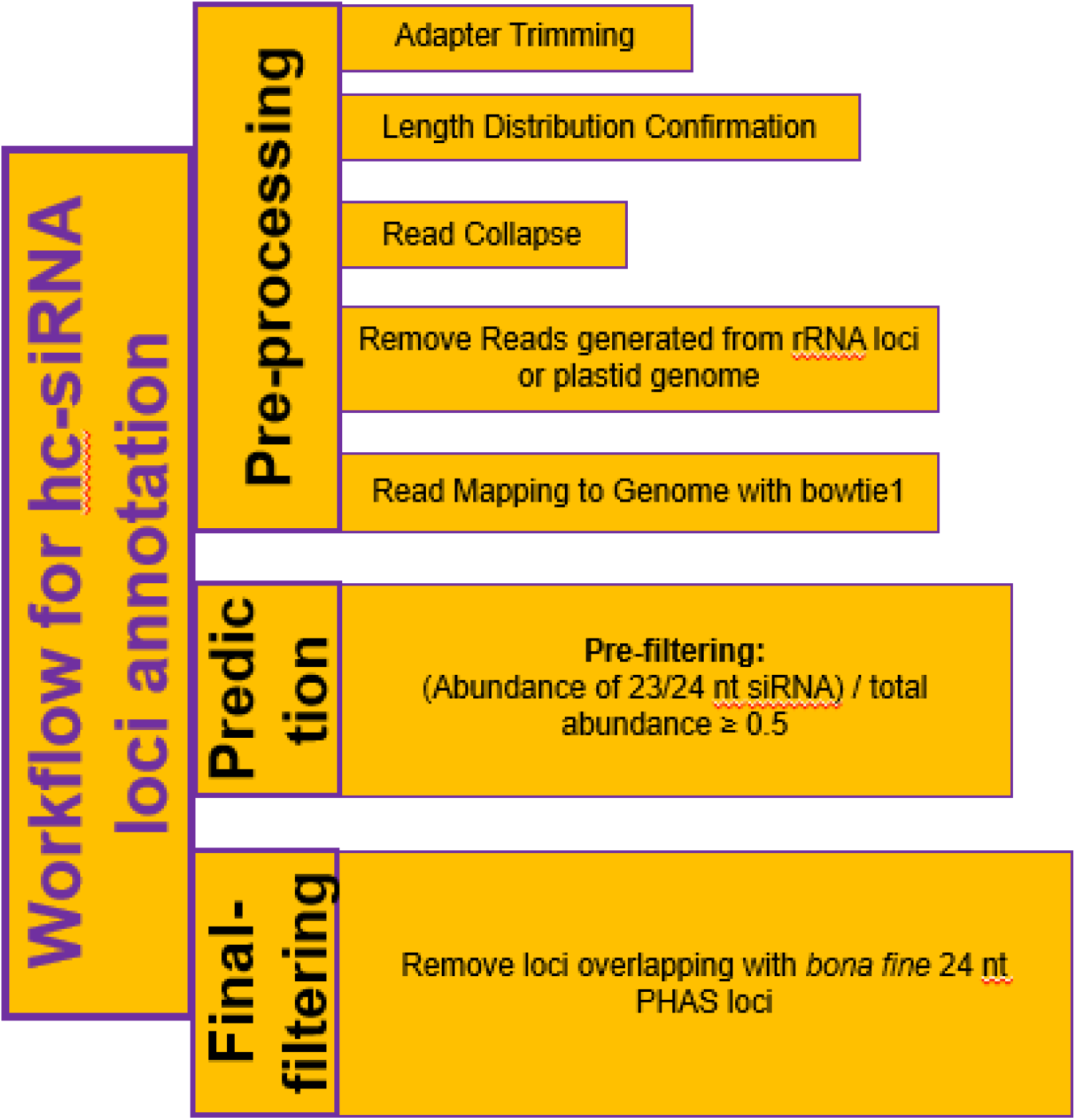
Workflow of hc-siRNA locus annotation.

**Figure S6.**
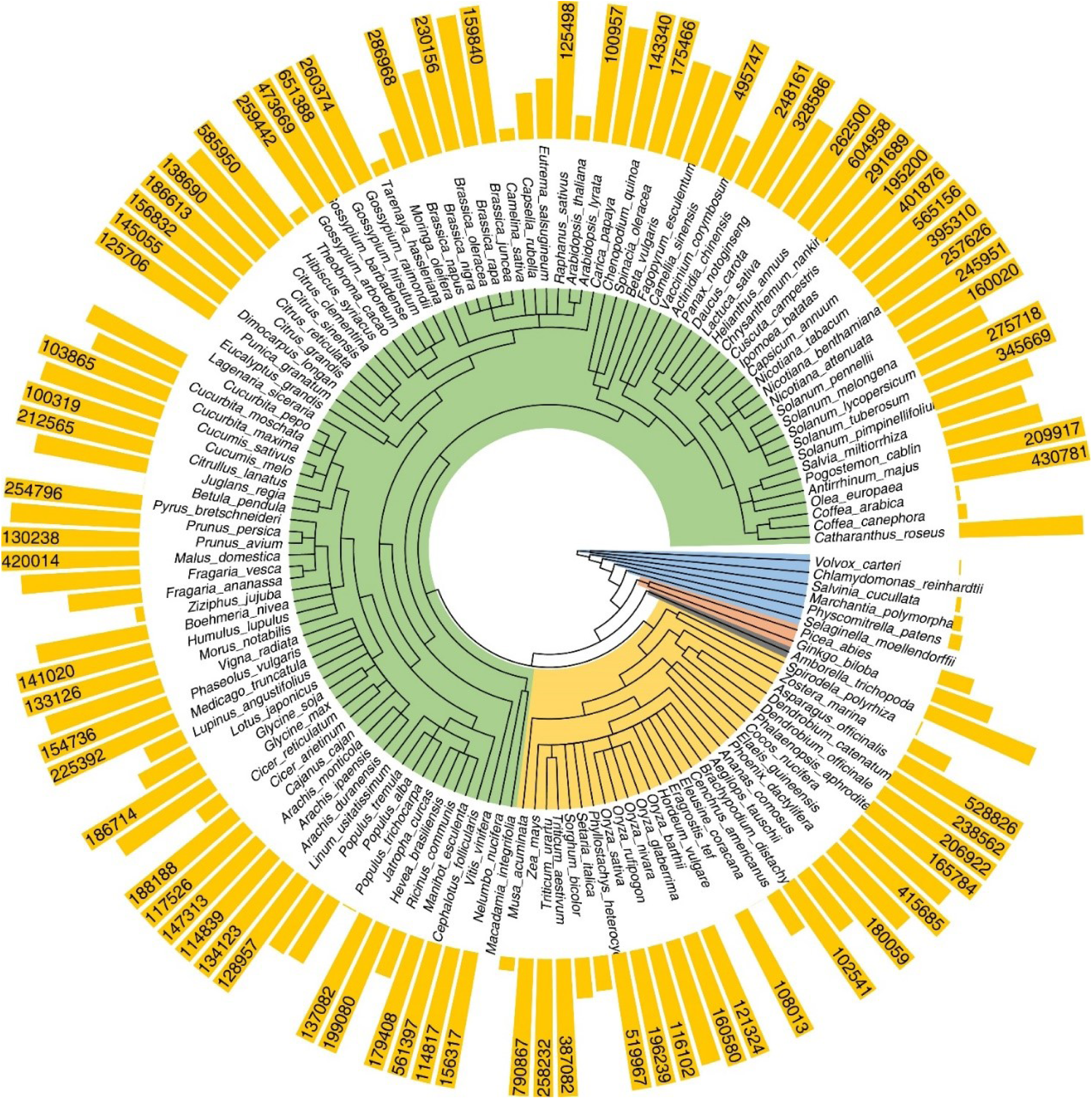
Overview of hc-siRNA loci of each species annotated in sRNAanno. Maximum value of bar plot is set to 100,000.

**Figure S7.**
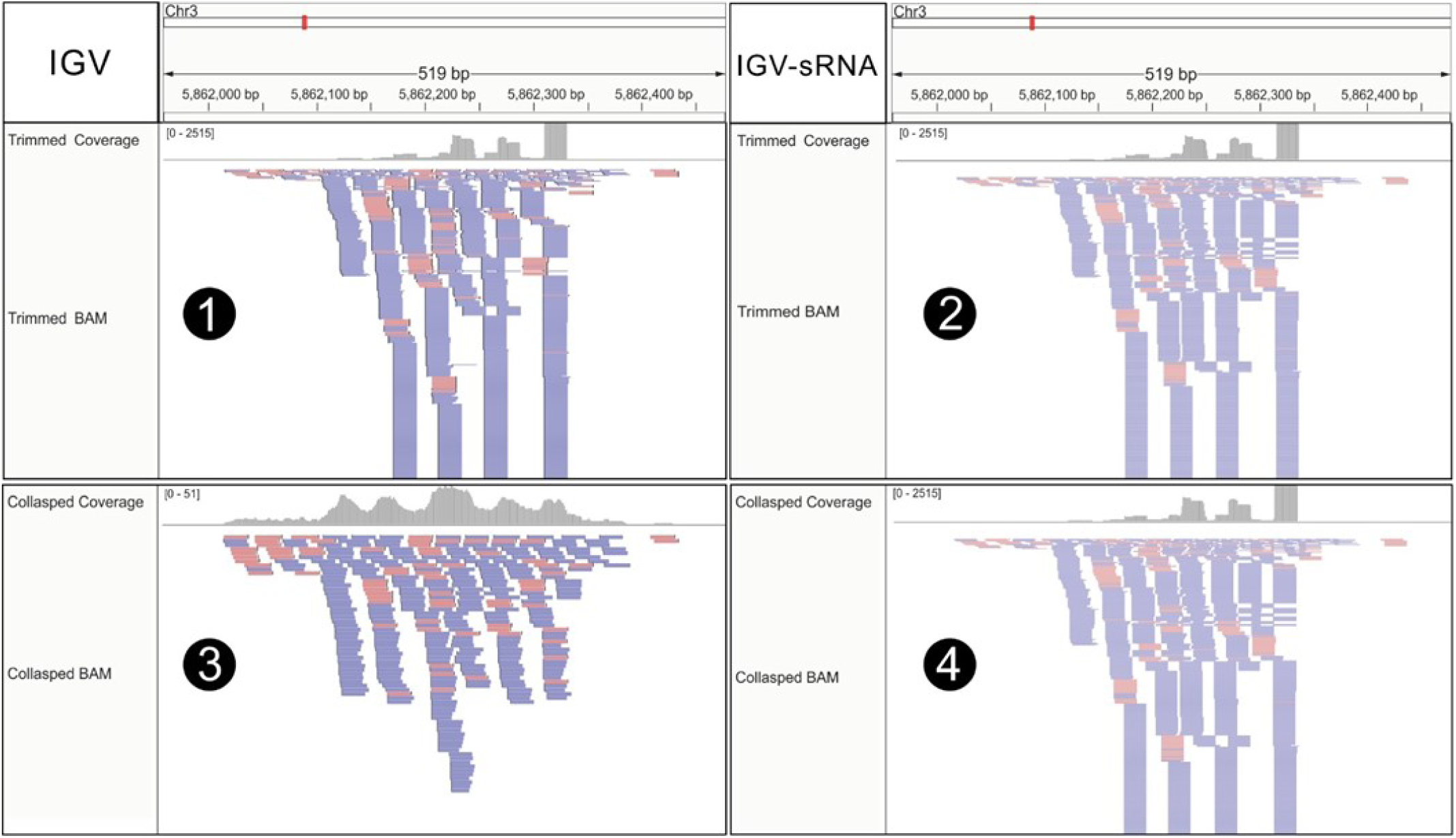
Comparison of IGV and IGV-sRNA on visualization of original (non-collapsed) and collapsed sRNA mapping files. (1) and (2) Alignments of IGV and IGV-sRNA are identical when they are viewing an original (non-collapsed) sRNA mapping file; (3) and (4) Alignments of IGV and IGV-sRNA are different when they are viewing a collapsed sRNA mapping file. The IGV shows each collapsed sRNA only once and the coverage values cannot be properly calculated.

## Notes

http://www.plantsRNAs.org

